# In Vitro Selection of Remdesivir-Resistant SARS-CoV-2 Demonstrates High Barrier to Resistance

**DOI:** 10.1101/2022.02.07.479493

**Authors:** Liva Checkmahomed, Venice Du Pont, Nicholas C. Riola, Jason K. Perry, Jiani Li, Danielle P. Porter, Guy Boivin

## Abstract

*In vitro* selection of remdesivir-resistant SARS-CoV-2 revealed the emergence of a V166L substitution, located outside of the polymerase active site of the nsp12 protein, after 9 passages. V166L remained the only nsp12 substitution after 17 passages at a final concentration of 10 µM RDV, conferring a 2.3-fold increase in EC_50_. When V166L was introduced into a recombinant SARS-CoV-2 virus, a 1.5-fold increase in EC_50_ was observed, indicating a high *in vitro* barrier to RDV resistance.

## Introduction

Remdesivir (RDV; VEKLURY^®^), a nucleotide prodrug inhibitor of the SARS-CoV-2 RNA-dependent RNA polymerase (nsp12) is one of the few approved antiviral drugs for the treatment of COVID-19 (1, 2). Characterization of clinical resistance development to this compound is currently in progress. So far, *in vitro* resistance to this antiviral has been rarely reported in SARS-CoV-2 and other viruses (3, 4).

In this study, we sought to select an RDV-resistant virus by serial passaging of a wild-type (WT) SARS-CoV-2 Canadian isolate in the presence of increasing concentrations of RDV in cell culture.

## Materials and methods

A WT SARS-CoV-2 isolate (D614G Spike + P323L nsp12 changes compared to WA1 strain) was passaged in Vero E6 cells initially in the presence of 1 µM of RDV and then increasing the drug concentration until clear virally-induced cytopathic effects were noted. A control virus was passaged concomitantly in the absence of drug during all the selection procedure. At specified RDV concentrations, 50% effective concentrations (EC_50_) values were determined by a plaque reduction assay (5) and the nsp12 genotype was determined by PCR amplification followed by Sanger sequencing.

Each non-silent nsp12 substitution was introduced into a recombinant SARS-CoV-2-Nano luciferase (Nluc) reporter virus. The reverse genetics system and the procedures required to assemble and produce SARS-CoV-2 infectious strains have been previously described in detail by Xie et al. (6). Antiviral activity (EC_50_ value) of RDV against SARS-CoV-2-Nluc was evaluated with a luciferase assay as reported (7) with some modifications.

Briefly, A549-hACE2 cells were seeded at 12,000 cells per well in medium containing 2% FBS into a white clear-bottom 96-well plate (Corning) at a volume of 50 µL and incubated overnight at 37°C with 5% CO_2_. On the following day, compounds were added directly to cultures as 3-fold serial dilutions with a Tecan D300e digital liquid dispenser, with DMSO volumes normalized to that of the highest compound concentration (final DMSO concentration <0.1%). SARS-CoV-2-Nluc viruses were diluted to MOI 0.05 and aliquoted 50 μL/well. After 48 h post compound treatment and infection, 75 μL of Nano luciferase substrate solution (Promega) was added to each well. Luciferase signals were measured using an Envision microplate reader (Perkin Elmer). The relative luciferase signals were calculated by normalizing the luciferase signals of the RDV-treated groups to that of the DMSO-treated groups (set as 100%).

A structural model of the SARS-CoV-2 polymerase complex with pre-incorporated RDV triphosphate (RDV-TP) was generated from the cryo-EM structure 6XEZ (8) as described elsewhere (9). V166 was substituted to an L, and all residues within 5 Å of this residue were conformationally sampled and minimized for both WT and V166L (10).

A total of N=6,920,015 genome sequences were deposited in GISAID database as of January 12, 2022. The tabulated amino acid substitutions from Wuhan-Hu-1 reference sequence (NC_045512) for each SARS-CoV-2 sequence (variant surveillance package) were downloaded through GISAID web portal (https://www.gisaid.org/) to assess the prevalence of the mutation.

## Results

The V166L substitution located in the nsp12 protein was detected after 9 passages reaching 6 µM of RDV (Table 1). This mutation was not present in the WT virus passaged in parallel in the absence of drug. The same substitution was detected at passages 11 and 17 (in presence of 8 and 10 µM of RDV, respectively). Further passages beyond 10 µM of RDV resulted in tiny plaques or absence of cytopathic effects. Virus isolates from passages 6, 10, and 17 exhibited a 3.9-, 2.7-, and 2.3-fold reduced susceptibility to RDV compared to the WT isolate using a plaque reduction assay in Vero E6 cells. The V166L substitution remained the only nsp12 mutation that emerged in the presence of RDV throughout the entire resistance selection process.

**Table 1:** Summary of cell culture passages for the selection of remdesivir-resistant SARS-CoV-2 virus with phenotypic and genotypic results.

| Passages | Remdesivir concentrations ($\mu\text{M}$) | EC<sub>50</sub> values (Fold change) | Non silent substitution in nsp12 |
| --- | --- | --- | --- |
| 0 | 0 $\mu\text{M}$ | 1.35 ± 1.00 $\mu\text{M}$ (1.0) | None |
| 1 | 1 $\mu\text{M}$ | | |
| 2 | 2 $\mu\text{M}$ | | |
| 3 | 4 $\mu\text{M}$ | | |
| 4 | 4 $\mu\text{M}$ | | |
| 5 | 5 $\mu\text{M}$ | | |
| 6 | 6 $\mu\text{M}$ | | |
| 7 | 6 $\mu\text{M}$ | | |
| 8 | 6 $\mu\text{M}$ | | |
| 9 | 6 $\mu\text{M}$ | 5.27 $\mu\text{M}$ (3.9) | V166L |
| 10 | 8 $\mu\text{M}$ | | |
| 11 | 8 $\mu\text{M}$ | 3.75 $\mu\text{M}$ (2.7) | V166L |
| 12 | 9 $\mu\text{M}$ | | |
| 13 | 9 $\mu\text{M}$ | | |
| 14 | 9 $\mu\text{M}$ | | |
| 15 | 9 $\mu\text{M}$ | | |
| 16 | 9 $\mu\text{M}$ | | |
| 17 | 10 $\mu\text{M}$ | 3.08 ± 0.95 $\mu\text{M}$ (2.3) | V166L |

**Table 1:** Remdesivir EC_50_ against SARS-CoV-2-Nluc WA1 reference and mutant viruses.

| Nsp12 substitution | RDV EC<sub>50</sub> (nM) |  |  | EC<sub>50</sub> fold-change from reference |
| --- | --- | --- | --- | --- |
| | 1<sup>st</sup> replicate | 2<sup>nd</sup> replicate | Average ± standard deviation | |
| WA1 WT reference | 49 | 65 | 57 ± 11 | 1.00 |
| P323L | 62 | 85 | 74 ± 16 | 1.30 |
| V166L+P323L | 88 | 89 | 88 ± 1 | 1.54 |

When introduced into a SARS-CoV-2-Nluc recombinant WT WA1 virus, the nsp12 P323L substitution (present at baseline in the Canadian strain) plus the V166L nsp12 substitution (selected with RDV) minimally increased the RDV EC_50_ value by 1.5-fold relative to the WT reference virus using a luciferase assay (EC_50_ P323L + V166L = 88 +/-1 nM; WT = 57 +/-11 nM, Table 2). A structural analysis identified residue 166 as located outside of the polymerase active site (Figure 1), but contacts residues in the active site Motifs A and D. Modeling suggests V166L provides only a modest perturbation of Motif D, therefore any impact from this substitution on RDV potency is suspected to be indirect. Furthermore, an extensive search in sequencing databases in January 2022 identified the V166L substitution in 0.004% (251/6920015) of SARS-CoV-2 sequences reported.

**Figure 1:**
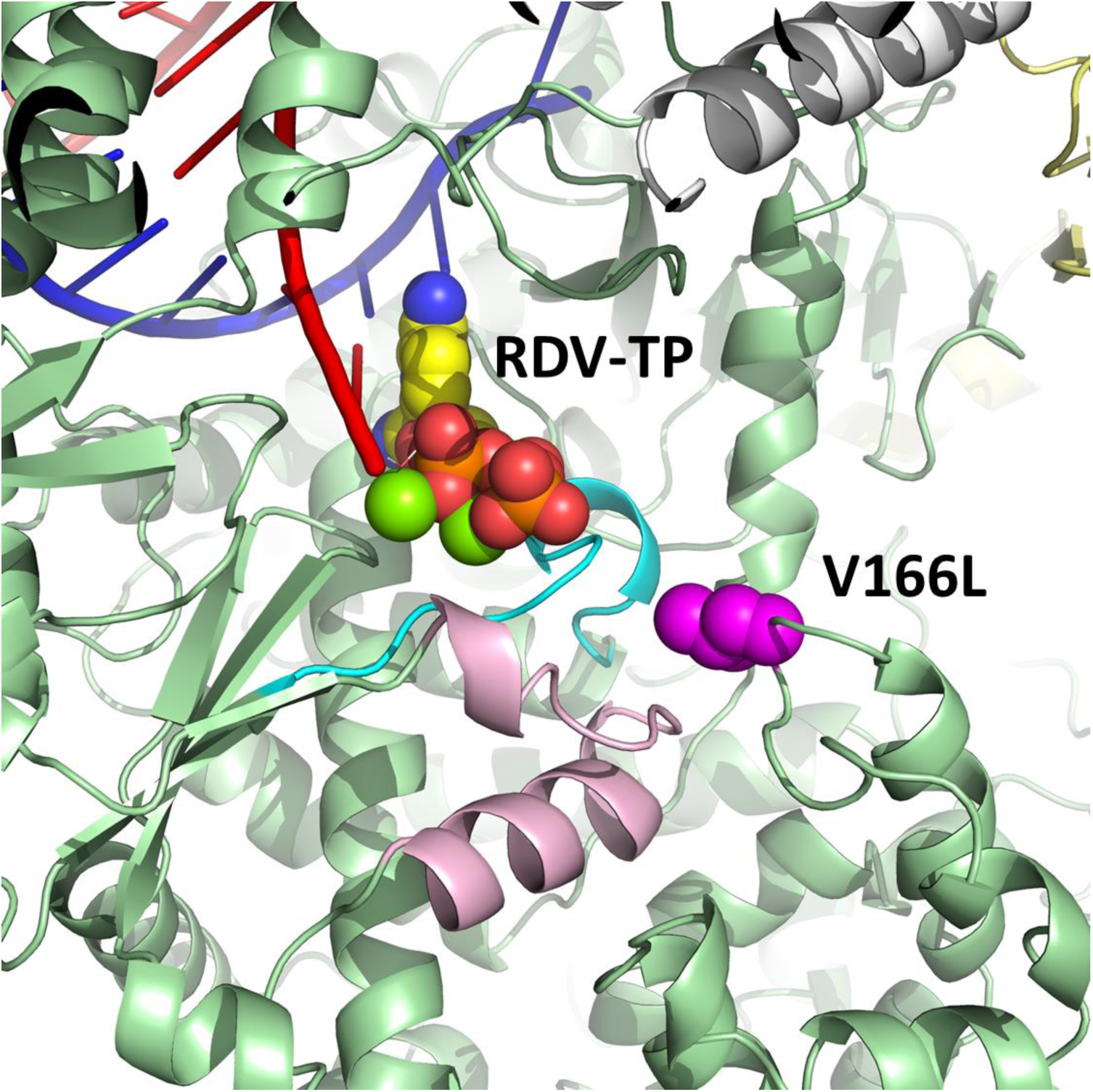
Model of pre-incorporated remdesivir triphosphate (RDV-TP) with a V166L substitution (magenta) in SARS-CoV-2 nsp12 (green). V166L is outside of the active site but contacts Motif A (cyan) and Motif D (pink) residues. Nsp7 and nsp8 are visible in white and yellow, respectively. The nascent RNA strand is in red and the template strand is in blue.

## Discussion

RDV is a broad-spectrum RNA polymerase inhibitor of several viral families including *Coronaviridae, Filoviridae, Pneumoviridae* and *Paramyxoviridae* (11, 12). Furthermore, it is the first antiviral approved by the FDA for the treatment of hospitalized cases of COVID-19. RDV resistance has been rarely documented in clinic (3, 4). Passaging SARS-CoV-2 *in vitro* in the presence of increasing concentrations of RDV selected a single substitution (V166L) in the viral polymerase nsp12 protein, which was associated with a minimal 1.5-fold increase in the RDV EC_50_ value after introduction into a recombinant virus. The low level of resistance was similar between the recombinant V166L virus and the various passages of the drug-selected strains (i.e., 1.5 and 2.3-3.9-fold). Interestingly, in an immunocompromised patient with prolonged SARS-CoV-2 shedding, the V166L substitution in nsp12 was observed after treatment with two 10-day courses of RDV and two units of convalescent plasma (unknown phenotype) (13).

Other *in vitro* studies have been conducted with coronaviruses that identified substitutions associated with resistance to RDV. Passage of the murine hepatitis virus (MHV) in presence of GS-441524 (parent nucleoside analog of RDV) selected two substitutions (F476L and V553L) after 23 passages in the viral RNA polymerase that led to 5-fold reduced susceptibility to RDV. Introduction of the corresponding mutations into SARS-CoV resulted in 6-fold reduced susceptibility to RDV *in vitro* and attenuated pathogenesis in a mouse model (3). Recently, the E802D substitution in nsp12 conferring a 2.5-fold increase in EC_50_ value was selected after serial passages of SARS-CoV-2 in presence of RDV *in vitro*. Notably, this substitution was also associated with a fitness cost (4).

The high number of passages reached at which only a single V166L substitution emerged and the minimal change in susceptibility to RDV associated with this substitution highlight the high genetic barrier of resistance for RDV.

## References

1. Beigel JH, Tomashek KM, Dodd LE, Mehta AK, Zingman BS, Kalil AC, Hohmann E, Chu HY, Luetkemeyer A, Kline S, Lopez de Castilla D, Finberg RW, Dierberg K, Tapson V, Hsieh L, Patterson TF, Paredes R, Sweeney DA, Short WR, Touloumi G, Lye DC, Ohmagari N, Oh MD, Ruiz-Palacios GM, Benfield T, Fatkenheuer G, Kortepeter MG, Atmar RL, Creech CB, Lundgren J, Babiker AG, Pett S, Neaton JD, Burgess TH, Bonnett T, Green M, Makowski M, Osinusi A, Nayak S, Lane HC, Members A-SG. 2020. Remdesivir for the Treatment of Covid-19 - Final Report. N Engl J Med 383:1813–1826.

2. Hall MD, Anderson JM, Anderson A, Baker D, Bradner J, Brimacombe KR, Campbell EA, Corbett KS, Carter K, Cherry S, Chiang L, Cihlar T, de Wit E, Denison M, Disney M, Fletcher CV, Ford-Scheimer SL, Gotte M, Grossman AC, Hayden FG, Hazuda DJ, Lanteri CA, Marston H, Mesecar AD, Moore S, Nwankwo JO, O’Rear J, Painter G, Singh Saikatendu K, Schiffer CA, Sheahan TP, Shi PY, Smyth HD, Sofia MJ, Weetall M, Weller SK, Whitley R, Fauci AS, Austin CP, Collins FS, Conley AJ, Davis MI. 2021. Report of the National Institutes of Health SARS-CoV-2 Antiviral Therapeutics Summit. J Infect Dis 224:S1–S21.

3. Agostini ML, Andres EL, Sims AC, Graham RL, Sheahan TP, Lu X, Smith EC, Case JB, Feng JY, Jordan R, Ray AS, Cihlar T, Siegel D, Mackman RL, Clarke MO, Baric RS, Denison MR. 2018. Coronavirus Susceptibility to the Antiviral Remdesivir (GS-5734) Is Mediated by the Viral Polymerase and the Proofreading Exoribonuclease. mBio 9.

4. Szemiel AM, Merits A, Orton RJ, MacLean OA, Pinto RM, Wickenhagen A, Lieber G, Turnbull ML, Wang S, Furnon W, Suarez NM, Mair D, da Silva Filipe A, Willett BJ, Wilson SJ, Patel AH, Thomson EC, Palmarini M, Kohl A, Stewart ME. 2021. In vitro selection of Remdesivir resistance suggests evolutionary predictability of SARS-CoV-2. PLoS Pathog 17:e1009929.

5. Niesor EJ, Boivin G, Rheaume E, Shi R, Lavoie V, Goyette N, Picard ME, Perez A, Laghrissi-Thode F, Tardif JC. 2021. Inhibition of the 3CL Protease and SARS-CoV-2 Replication by Dalcetrapib. ACS Omega 6:16584–16591.

6. Xie X, Muruato A, Lokugamage KG, Narayanan K, Zhang X, Zou J, Liu J, Schindewolf C, Bopp NE, Aguilar PV, Plante KS, Weaver SC, Makino S, LeDuc JW, Menachery VD, Shi PY. 2020. An Infectious cDNA Clone of SARS-CoV-2. Cell Host Microbe 27:841–848 e3.

7. Xie X, Muruato AE, Zhang X, Lokugamage KG, Fontes-Garfias CR, Zou J, Liu J, Ren P, Balakrishnan M, Cihlar T, Tseng CK, Makino S, Menachery VD, Bilello JP, Shi PY. 2020. A nanoluciferase SARS-CoV-2 for rapid neutralization testing and screening of anti-infective drugs for COVID-19. Nat Commun 11:5214.

8. Chen J, Malone B, Llewellyn E, Grasso M, Shelton PMM, Olinares PDB, Maruthi K, Eng ET, Vatandaslar H, Chait BT, Kapoor TM, Darst SA, Campbell EA. 2020. Structural Basis for Helicase-Polymerase Coupling in the SARS-CoV-2 Replication-Transcription Complex. Cell 182:1560–1573 e13.

9. Gordon CJ, Lee HW, Tchesnokov EP, Perry JK, Feng JY, Bilello JP, Porter DP, Gotte M. 2021. Efficient incorporation and template-dependent polymerase inhibition are major determinants for the broad-spectrum antiviral activity of remdesivir. J Biol Chem doi:10.1016/j.jbc.2021.101529:101529.

10. Schrödinger Software Release 2021-3. 2021. Schrödinger, LLC, New York, NY.

11. Xu Y, Barauskas O, Kim C, Babusis D, Murakami E, Kornyeyev D, Lee G, Stepan G, Perron M, Bannister R, Schultz BE, Sakowicz R, Porter D, Cihlar T, Feng JY. 2021. Off-Target In Vitro Profiling Demonstrates that Remdesivir Is a Highly Selective Antiviral Agent. Antimicrob Agents Chemother 65.

12. Lo MK, Jordan R, Arvey A, Sudhamsu J, Shrivastava-Ranjan P, Hotard AL, Flint M, McMullan LK, Siegel D, Clarke MO, Mackman RL, Hui HC, Perron M, Ray AS, Cihlar T, Nichol ST, Spiropoulou CF. 2017. GS-5734 and its parent nucleoside analog inhibit Filo-, Pneumo-, and Paramyxoviruses. Sci Rep 7:43395.

13. Kemp SA, Collier DA, Datir RP, Ferreira I, Gayed S, Jahun A, Hosmillo M, Rees-Spear C, Mlcochova P, Lumb IU, Roberts DJ, Chandra A, Temperton N, Collaboration C-NBC-, Consortium C-GU, Sharrocks K, Blane E, Modis Y, Leigh KE, Briggs JAG, van Gils MJ, Smith KGC, Bradley JR, Smith C, Doffinger R, Ceron-Gutierrez L, Barcenas-Morales G, Pollock DD, Goldstein RA, Smielewska A, Skittrall JP, Gouliouris T, Goodfellow IG, Gkrania-Klotsas E, Illingworth CJR, McCoy LE, Gupta RK. 2021. SARS-CoV-2 evolution during treatment of chronic infection. Nature 592:277–282.

